# Energetic consequences of single and repeated freezing in the intertidal mussel, *Mytilus trossulus*

**DOI:** 10.1101/2025.02.10.637536

**Authors:** Josh Chia Chi Yang, Katie E. Marshall

## Abstract

The bay mussel, *Mytilus trossulus*, is a bivalve commonly found in intertidal zones along the west coast of North America which risks freezing during low tides in the winter. Previous work has shown that despite their freeze tolerance, freezing can still cause damage. Yet little is known about the energetic consequences of these freezing events. Here we measured the oxygen consumption of mussels before and after single and repeated freezing exposures across three seasons. We compared these responses to hypoxia exposures as tissues are not perfused while frozen which inhibits gas exchange. In general, we observed that mussels’ metabolic rates decreased immediately after a single freeze but recovers after 24 hours. By contrast, repeated freeze-thaws caused variable effects of metabolic rate, depending on the season of exposure. Overall, this suggests that the periods of recovery that occur between freeze-thaw cycles may mitigate freezing damage. We also found that hypoxia in general causes an increase in metabolic rate and may be associated with clearing oxygen debt. Therefore, the metabolic consequences after freezing are unlikely to be driven by hypoxia stress. Here we explore the energetic costs of freezing in *M. trossulus* informing future research directions into the mechanisms behind freeze tolerance.

## Introduction

Animals living in areas that experience sub-zero temperatures face the challenges of surviving low temperatures and the risk of freezing. This is especially relevant for ectotherms as they are largely physiologically unable to regulate their body temperatures and instead may rely on behaviourally avoiding the cold. However, in the intertidal zone, many ectotherms such as mussels and barnacles are largely sessile. Thus, they are particularly likely to experience freezing during the winter low tides. As such, many intertidal ectotherms in temperate and polar regions have developed the capability to survive freezing (Aarset, 1982; Gill et al., 2024).

To survive freezing, animals must also survive the associated physical, physiological, and biochemical challenges. One example of a physical stressor is damage due to ice formation during freezing inside the animal (Toxopeus & Sinclair, 2018). Another freezing-induced stressor is the disruption of the extracellular-intracellular ionic gradient, which can in turn impair the membrane potential necessary for neuromuscular function (Baust, 1973; Boardman et al., 2011; Zachariassen & Hammel, 1976). Freezing also induces hypoxia stress when tissues become ischemic due to the lack of circulation and gas exchange (Williams, 1970). Additional challenges emerge during thawing as the resumption of respiration and circulation cause the rapid re-oxygenation of cells which can lead to ischemia-reperfusion injury (Cadenas & Davies, 2000; J. M. Storey & Storey, 2005). Despite being able to survive these stressors during each freezing bout, these consequences of freezing may impose energetic consequences to the animal.

The energetic costs of freezing at the whole organism level in marine organisms have yet to be investigated. Coping with oxygen debt due to the periods of hypoxia or repair of damaged tissues may consume cellular ATP and potentially represent a metabolic cost (Ellington, 1983). In the freeze-tolerant mussel, *Geukensia demissus*, ATP and ADP were reduced following freezing (Storey & Churchill, 1995). Given the synthesis of ATP is a largely aerobic process, the associated costs may manifest at the whole-organism level as increases in oxygen consumption rate.

Alternatively, if freezing damages the oxygen cascade and impairs oxygen uptake, then the lowered capacity to supply aerobic metabolism with oxygen may result in a metabolic depression. For example, freezing can damage the gills of aquatic organisms, which may reduce their capacity for oxygen uptake and hinder aerobic metabolism (Kennedy, 2022). For example, CO_2_ production in the flightless moth *Pringleophaga marioni* decreases immediately after freezing (Sinclair et al., 2004). Thus, it is also possible that damage incurred during freezing may limit aerobic metabolism and result in a metabolic depression.

However, a depression in metabolic rate may also be a result of active suppressive mechanisms. Sessile intertidal animals are equipped with the ability to downregulate metabolism (K. B. Storey, 1988) and this has been observed as a response to thermal stress (Hui et al., 2020). While there are no studies in intertidal animals investigating metabolic rates in response to low temperatures to date, reduction in metabolic rates have been observed in other terrestrial invertebrates post-thawing (Sinclair et al., 2004, 2013).

Typically, studies regarding freeze tolerance only address a single freeze-thaw cycle, but multiple freezing events can occur in the intertidal during a prolonged cold spell due to tidal cycles (Kennedy et al., 2020). Multiple freeze-thaw cycles may yield different physiological consequences compared to a single freeze-thaw cycle. *Mytilus trossulus* undergoing multiple freeze-thaw cycles have a higher survival rate than a single prolonged freeze with freezing time held constant (Gill et al., 2023) while the woolly bear caterpillar *Pyrrharctia isabella* demonstrates the opposite (Marshall & Sinclair, 2011). It is possible that mussels are more tolerant to stress resulting from repeated freeze-thaw cycles or that they are more capable of repair following freezing damage. For example, the gills of *M. trossulus* after freezing appear to repair over time (Kennedy, 2022). As these repair processes likely cost ATP, periods between freezing may serve as a temporal buffer and provide an opportunity to regenerate the consumed ATP to fuel repair. Thus, repeated freeze-thaw cycles (with total freezing time held constant) may produce relatively less metabolic perturbation than single freezing events due to periods of interbout repair.

In many species, freeze tolerance is either only present or most robust in the winter months as preparation for freezing begins in the late summer and increases until winter (Lee, 2010; Marshall et al., 2014; Murphy & Johnson, 1980). In *M. trossulus*, freeze tolerance is present year round and is higher in the winter than in the summer (Kennedy et al., 2020). This may be in part driven by increasing winter salinities at that particular site but it may also be partially supported by a reduction of basal metabolic rates in the winter found in *M. edulis* (Hatcher et al., 1997) may dampen the negative effects of ischemia-reperfusion during thawing. Therefore, the overall energetic cost of freezing may be relatively lower in the winter due to increased freeze tolerance and lower metabolic rates.

In this study we use the bay mussel, *M. trossulus*, to examine the energetic costs of freezing. This species inhabits the intertidal zones along the west coast of North America (Hilbish et al., 2000), and is relatively sessile and cannot behaviorally avoid sub-zero temperatures making them an ideal model for freeze tolerance research. Our study site at Tower Beach in Vancouver, B.C., Canada experiences mixed semidiurnal tides, with the lowest low tide occurring at night during the winters when air temperatures are the lowest (Kennedy et al., 2020). Therefore, mussels in this area are exposed to freezing when air temperatures are sufficiently low and thus serve as a viable model for this study.

The objective of this study is to investigate the whole organism metabolic rate of mussels before and after a freezing event and whether this is affected by season and by frequency of exposure. If freezing decreases metabolic rate due to damage to the oxygen cascade, then we predict that oxygen consumption would decrease after freezing. Conversely, if freezing increases metabolic rate due to restorative processes such as ATP synthesis, then we predict that oxygen consumption will increase after freezing. We also hypothesize that freezing in the winter will be the least damaging due to winter acclimation and therefore we predict that oxygen consumption rates will be relatively less perturbed following freezing in the winter. Next, we also hypothesize that freezing repeatedly will be less energetically costly due to periods of recovery in between each freeze and we predict that oxygen consumption will be lower after repeated freezes than single freezes. Using closed system respirometry, we measure whole organism oxygen consumption rate as a proxy for metabolic rate to compare aerobic metabolism before and after freezing. This study provides insight on the energetic costs of both single and repeated freezing events and whether this cost differs across seasons in an intertidal mussel.

## Materials and Methods

### Field Collection and Laboratory Acclimation

We collected all mussels in our experiments from Tower Beach, Vancouver, B.C., Canada (49°16’26.1∝N 123°15’23.7”W). The tides in this area are mixed semi-diurnal, with two low and two high tides of different magnitudes daily. Mussel collections were completed under a Scientific Licence, Management of Contaminated Fisheries Regulations from the Department of Fisheries and Oceans Canada (Licence numbers: XMCFR 34 2021, XMCFR 34 2022). We measured air temperatures, as well as salinity and water temperatures at approximately 25-50 cm water depth with a YSI handheld conductivity meter (Pro 30 series with a PRO 30 COND-T probe, Xylem, Yellow Springs, Ohio). Mussels were collected from the same mussel bed located directly in front of the gun tower at Tower Beach from the mid-intertidal zone on each sampling day.

We selected mussels with shell lengths of 2-4 cm as larger mussels would be unable to fit into the experimental equipment. After collection, mussels were transported to 20L aquaria filled with seawater (20 ppt). Natural seawater was sourced from Vancouver Aquarium and mixed with dechlorinated water to adjust to the desired salinity. The aquaria were placed in incubators which maintained a 12:12 light:dark cycle and held the temperature at 15 °C (MIR-154, Sanyo, Bensenville, USA). Tank water was aerated using air stones and changed every 2-3 days. Epibionts were removed and mussels were haphazardly assigned to experimental groups. Mussels were fasted and used 2-11 days after the date of collection. Prior to experimentation, mussel shell lengths were measured anterior to posterior at the longest length using digital calipers (VC-HeiKa, FineSource).

### Experimental Temperature Exposures

For experimental exposures, mussels were removed from the aquarium and placed individually in 25 mm *Drosophila* tubes. Then, a 24 AWG copper-constantan type-T thermocouple (OMEGA Engineering, St-Eustache, Quebec) connected to Picolog TC-08 interfaces (Pico Technology, Cambridge, UK) was attached to the shell of each mussel and secured using cotton or styrofoam. Temperature data was collected using PicoLog 6 beta software to track mussel body temperatures and determine freezing events. This was demonstrated as a rapid release of heat and dramatic spike in body temperature caused by ice formation (i.e. the supercooling point demonstrated in Fig. S.1; Lee, 2010). Mussels were placed into wells in an aluminum head (insulated by foam) that was cooled by a circulating bath with approximately 1:1 methanol and water mixture (Thermofisher, PC200 Model or Lauda, ECO Silver: RE 415 S Model, Wurzburg, Germany). Each individual mussel only underwent one temperature exposure.

To conduct experimental exposures, the circulator was initially set at the acclimation temperature of 15 °C and cooled at −1.25 °C/min until the experimental temperature. This cooling rate reflects the rapid cooling that *M. trossulus* would experience in the field when the tide recedes and they are suddenly exposed sub-zero air temperatures (Kennedy et al., 2020). The freezing temperature was chosen to be −6.2 °C as pilot trials ensured 100% of mussels would freeze within 2 hours when exposed to this temperature.

For single freezing exposures, mussels were exposed for 6 hours at −6.2 °C. For repeated freezing exposures, mussels were exposed for 2 hours at −6.2 °C for 3 consecutive days with 22 hours between each freezing exposure. For hypoxia exposures, the mussels were instead held at 15 °C for the duration of the exposures, mimicking emersion during a low tide. These exposures were conducted at the same time of day as the respective freezing exposures. To test the effects of supercooling on oxygen consumption, mussels were exposed to 6 hours at their supercooling point −5.5 °C (Kennedy et al., 2020). Summary of experimental exposures and environmental measurements throughout the study are available in Table S1. All exposures were sublethal as no mortality was observed in the week following an experimental exposure.

### Closed Respirometry

Mussels were placed into individual 80 mL glass chambers filled with fully aerated seawater (20 ppt) and sealed with rubber stoppers. Temperature was maintained at 15 °C during each experiment using an incubator (MIR-154, Sanyo, Bensenville, USA). Chamber water was mixed using a magnetic stir bar and mussels were elevated to prevent contact with the stir bar. Oxygen content was measured with Oxy-4 SMA trace (G3), Polymer Optical Fibers, and Oxygen Sensor Spot SP-PSt3-NAU and the water temperature was taken using a Pt100 Temperature Sensor (PreSens Precision Sensing, Regensburg, Germany), and recorded using PreSens Measurement Studio 2.0 Software. Atmospheric pressure was also recorded by the Oxy-4 SMA trace (G3). The sampling frequency was 0.33 Hz. Oxygen sensors were calibrated to 0% oxygen saturation with continuous bubbling of nitrogen gas and 100% oxygen saturation with aerated seawater using air stones.

Oxygen consumption of mussels was measured for one hour prior and immediately after experimental exposures for at least 60 minutes. Specifically for repeated exposures, the final oxygen consumption rate was measured after the final exposure. For the single and repeated exposures in the winter and the summer, an additional measurement was conducted after 24 hours of recovery at 15 °C. The volume of water after each trial was measured and adjusted for 9 in the analysis. Background oxygen consumption for each trial was measured in a separate empty chamber and subtracted from mussel oxygen consumption rates in the analysis.

### Respirometry Data Processing

We determined the first moment of detectable reduction in oxygen content from the raw oxygen content traces, defined by deviance from the background respiration rate. Chambers which did not display a clear signal of detectable reduction in oxygen content were excluded. This occurred for one mussel in the summer repeated hypoxia and summer single freeze treatments (Fig. S.2). Then, we sectioned the raw oxygen content values into four 10 minute windows 10 minutes after this first moment of detectable reduction in oxygen content. The slope for each section, representing the rate of oxygen consumption, was calculated using the “respR” package (Harianto et al., 2019) The mean across the 40 minute time period was then calculated for each individual and used as its oxygen consumption rate. The background was adjusted by subtracting it from the calculated oxygen consumption rate. Measurements were normalized to the volume of seawater in the respirometer measured after each trial. Oxygen consumption rate in this study is expressed in a mass-independent way (µg O_2_/h) as mussel mass measured by proxy with shell length across treatment groups were not significantly different.

### Statistical Methods

All statistical analyses were performed using R (v. 4.2.1; R Development Core Team, 2022). To compare the oxygen consumption rates before and after freezing or hypoxia exposure of the same individual, we used a linear mixed effects model with the individual as the random effect (MO_2_ ∼ timepoint + (1|individual)) and used a Type 2 analysis of deviance to evaluate significance. Linear mixed effect models were built using the “lme4” package (Bates et al., 2015). To compare the effects among treatment (freezing and hypoxia), frequency (single and repeated exposures), and season (fall, winter, and summer), a ratio was calculated by dividing the post-exposure MO_2_ by the mussel’s initial pre-exposure baseline MO_2_. Then, to compare MO_2_ ratios across seasons, frequency, and treatment, we used an ANOVA on a linear model (MO_2_ ratio∼ season*treatment*frequency). To test the effects of supercooling on MO_2_ ratios, mussels were separated based on outcome after a 6 hour −5.5 °C exposure into either frozen or supercooled (not frozen). ANOVA was used to compare MO_2_ ratios across mussels frozen when exposed to −6.2 °C, mussels exposed to −5.5 °C but were supercooled, and mussels exposed to - 5.5 °C but frozen. To test the effects of recovery, another ratio was calculated by dividing the MO_2_ measured 24-hours post-treatment the pre-treatment MO_2_. We tested for the effect of recovery time using a linear mixed effects model with individual as the mixed effect (MO_2_ ∼ hours post treatment + (1|individual) and evaluated the significance with a type 2 analysis of deviance.

## Results

As mussels were sampled across three seasons, we first wanted to examine whether their routine metabolic rate (i.e. before freezing or hypoxia exposure) varied across seasons. We compared the baseline MO_2_ of all mussels (i.e. MO_2_ before exposures) which were included in the subsequent analyses. We found that MO_2_ did not significantly vary among seasons (ANOVA; F_2,93_=0.24, p=0.79; Fig. 1). One mussel from the single freeze exposure and the single hypoxia exposure were excluded from the summer cohort as there was no detectable oxygen consumption during the measurement window.

**Figure 1.**
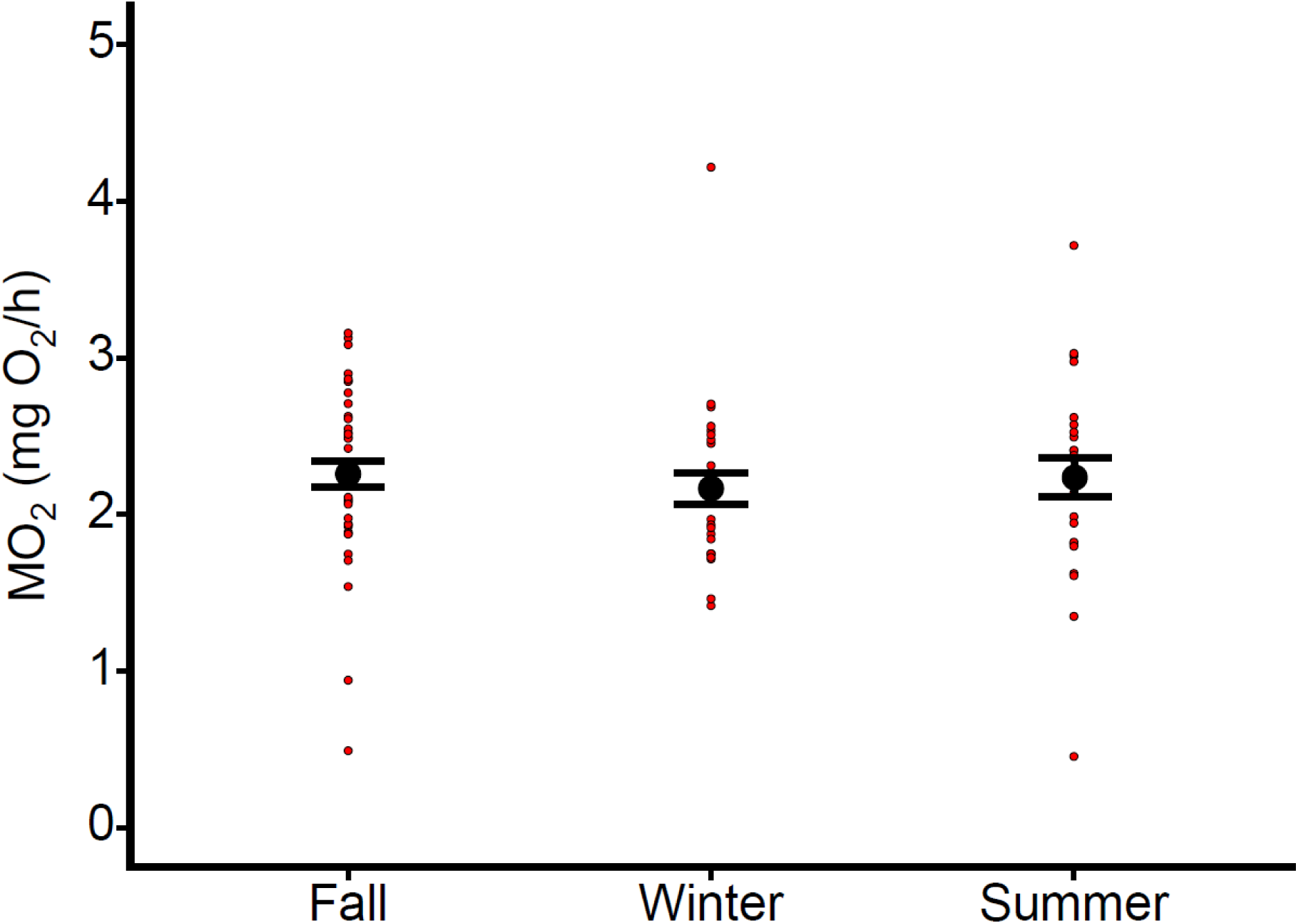
Seasonal variation in *M. trossulus* routine MO_2_. Comparison of the mean routine MO_2_ measured in water at 15 °C of *M. trossulus* collected intertidally across fall (n=42), winter (n=28), and summer (n=26). Individual MO_2_ values were calculated based on the mean MO_2_ over a 40 minute period of detectable oxygen consumption immediately before experimental exposures. Large black points represent the mean, errors bars represent the standard error of the mean, and smaller red points represent the individual data points.

**Figure 2.**
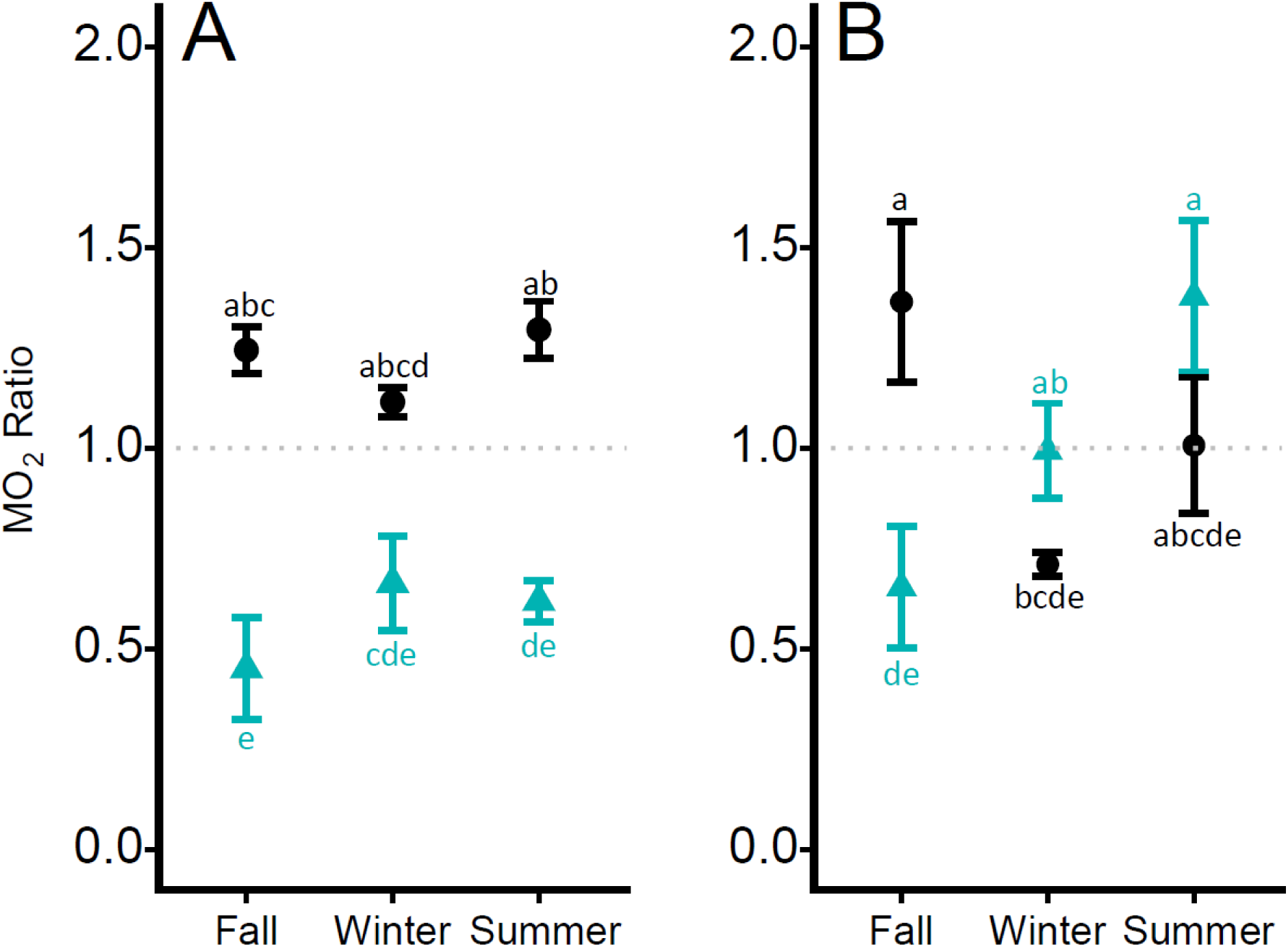
Mean MO_2_ Ratio of *M. trossulus* across seasons, treatment, and frequency. Mean ratio of MO_2_ of *M. trossulus* immediately after exposure to freezing or hypoxia singly (A) or repeatedly (B). Ratios were obtained by dividing MO_2_ obtained after exposure to pre-exposure MO_2_. A ratio above 1 (grey dotted line) generally signifies an increase in MO_2_ and a ratio below 1 generally signifies a decrease in MO_2_. For each treatment, n=7 except for single freeze and repeated hypoxia in the summer which had n=6. Teal triangles represent mean MO_2_ ratios after freezing, and black circles represent the mean MO_2_ ratios after hypoxia exposure. Error bars signify the standard error of the mean. Letters display Tukey post hoc results across season, frequency, and treatment.

We then subjected the mussels either to freezing or hypoxia exposure and examined the effects on MO_2_ immediately after exposure. For freezing exposures, mussels were exposed to - 6.2 °C and internal ice formation were confirmed by the presence of a supercooling point (Fig. S.1). Hypoxia exposures were conducted at 15 °C, which was the maintenance temperature. The mean MO_2_ over a 40 minute period was calculated for mussels before and after experimental exposures (Fig. S.3). The MO_2_ before and after treatment were then compared using a repeated measures ANOVA with the individual as the random effect. We found that after a single freeze in all seasons, there is always a significant decrease in MO_2_ (Fig. S.3 A-C). After repeated freezes, mussels in the fall showed a significant decrease in MO_2_ (Fig. S.3 G), mussels in the winter did not show statistically different MO_2_ after treatment (Fig. S.3 H), and mussels in the summer showed a significant increase in MO_2_ (Fig. S.3 I). After a single hypoxia exposure, there is a significant increase in MO_2_ in all seasons (Fig. S.3 D-F). After repeated hypoxia exposures, mussels in the fall and the summer showed a significant increase in MO_2_ (Fig. S.3 J & L respectively) while mussels in the winter showed a significant decrease in MO_2_ (Fig. S.3 K). Statistical results are reported in Table S2.

To control for individual variation in MO_2_, we normalized each individual’s post-exposure MO_2_ to their pre-exposure MO_2_ (baseline MO_2_) and calculated a ratio. Then we compared across groups using these ratios. To compare the effects of season and frequency (i.e. single vs repeated) between freezing and hypoxia on mussel MO_2_, we conducted a three-way ANOVA. The three-way interaction between treatment, season, and frequency was significant (F_2,70_=3.89, p=0.025) where the effect of repeated exposures depends on season and treatment. There are also significant two-way interactions between season and treatment (F_2,70_=8.85, p<0.001) as well as treatment and frequency (F_1,70_=18.00, p<0.001) on MO_2_ ratio. Treatment had a significant effect on MO_2_ ratio (F_1,70_=21.64, p<0.001) where MO_2_ ratio was generally elevated after hypoxia exposures and depressed after freezing. Season was nearly significant (F_2,70_=3.04, p=0.054) and likely driven by the lower MO_2_ ratio in the fall.

To test whether crossing the freezing threshold was a primary driver for metabolic changes after freezing, mussels were exposed to the average supercooling point (−5.5 °C) determined in Kennedy et al 2020. When exposed to the average mussel supercooling point for 6 hours, 5/14 (36%) mussels froze and the 9/14 (64%) did not. Using a repeated measures ANOVA with the individual as the random effect, we compared the MO_2_ before and after exposures. For mussels which froze, MO_2_ ratio after cold exposure was significantly lower than before cold exposure (*χ*^2^=23.59, df=1, p<0.0001). For mussels which did not freeze after exposure to −5.5 °C, the MO_2_ was not significantly different relative to the baseline (*χ*^2^=0.0011, df=1, p=0.97).

Similarly, a ratio was calculated and we compared between mussels frozen at −6.2 °C, mussels which did not freeze at −5.5 °C, and mussels which froze at −5.5 °C. We found no significant differences among these groups (ANOVA; F_2,18_=2.7986, p=0.087; Fig. 3). However, we noticed a trend where mussels were frozen at −5.5 °C and −6.2 °C showed similar MO_2_ ratios, and that mussels which did not freeze but exposed to −5.5 °C or 15 °C showed similar MO_2_ ratios.

**Figure 3.**
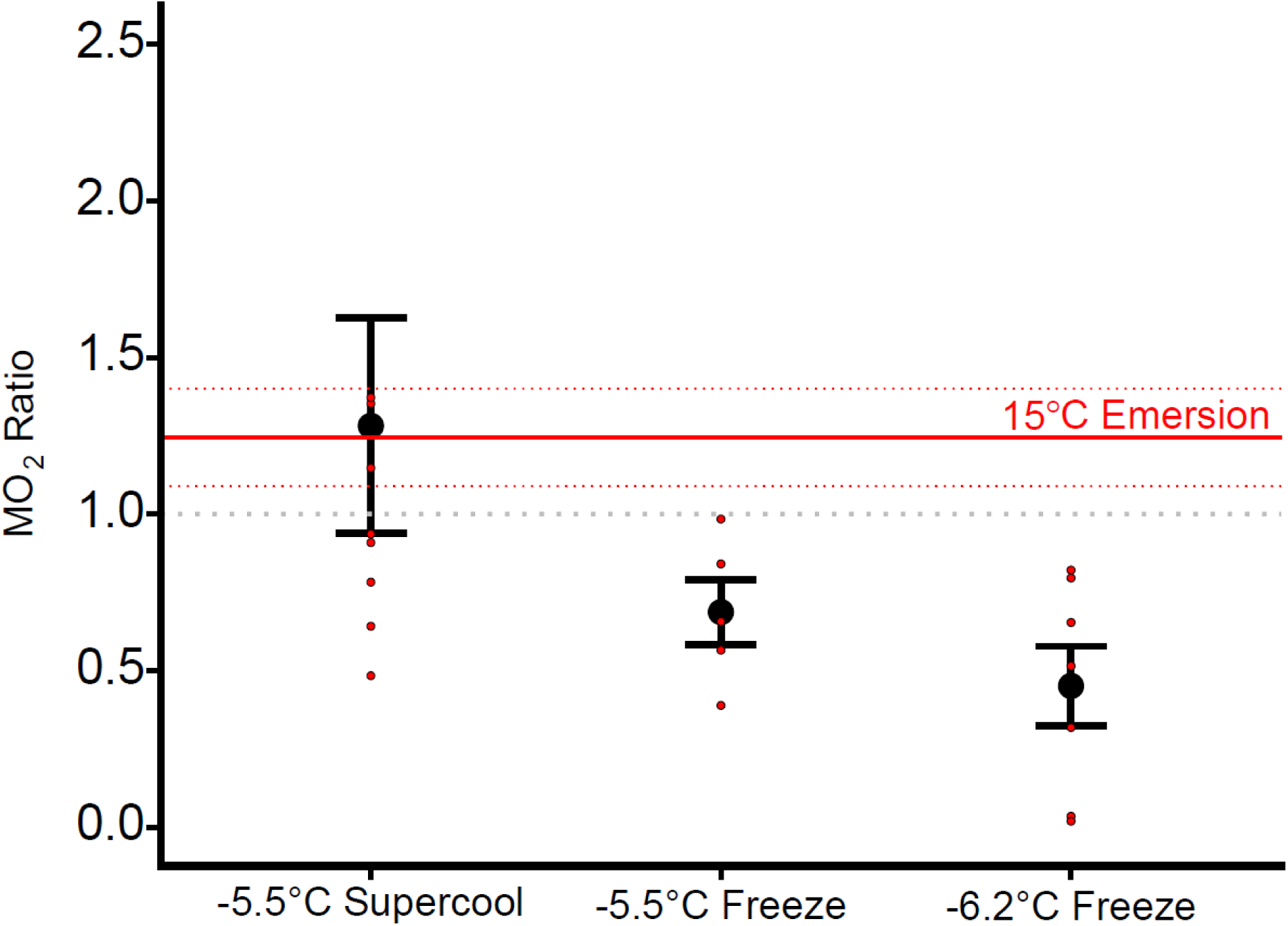
Effects of supercooling on MO_2_. Mean MO_2_ ratio of *M. trossulus* immediately after a single exposure to either −5.5 °C (i.e. supercooling point) or −6.2 °C. At the supercooling point, 5/14 mussels froze while 9/14 did not and their MO_2_ were separated based on the outcome. All mussels froze at 6.2 °C. The mean MO_2_ ratio of *M. trossulus* immediately after a hypoxia exposure is indicated by a red solid line ± SEM (red dotted line). Ratios were obtained by normalizing the MO_2_ after exposure to the baseline. A ratio above 1 (grey dotted line) generally signifies an increase in MO_2_ and a ratio below 1 generally signifies a decrease in MO_2_. n=7 for the single freeze at –6.2 °C and for the single hypoxia. Large black points represent the mean, error bars signify the standard error of the mean, and smaller red points represent the individual data points.

**Figure 4.**
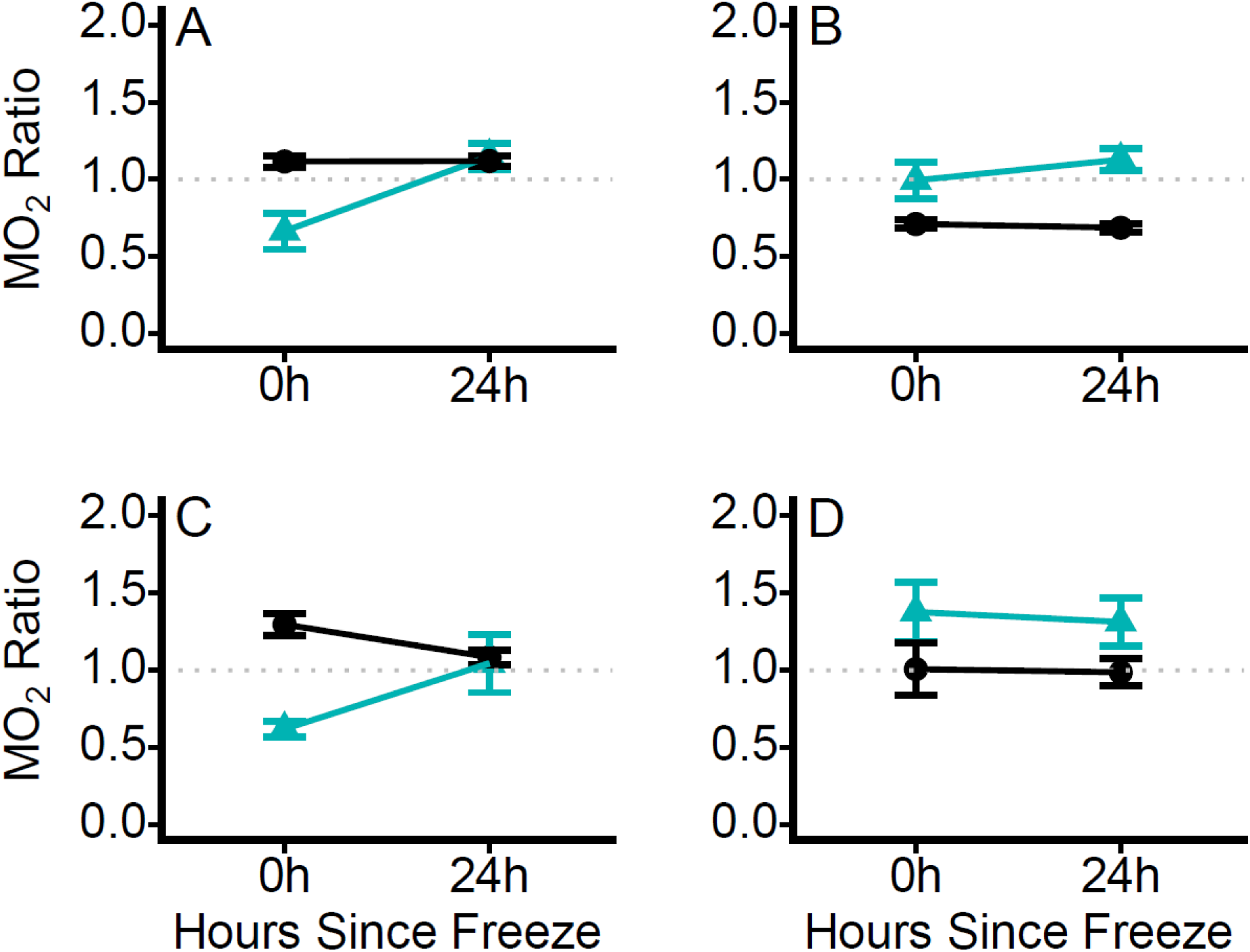
Mean MO_2_ ratio of *M. trossulus* immediately and 24 hours after single or repeated freeze or hypoxia exposures. Mussel MO_2_ was measured immediately and 24 hours after exposure to (A) single freeze and hypoxia exposures in the winter, (B) repeated freeze and hypoxia exposures in the winter, (C) single freeze and hypoxia exposures in the summer, and (D) repeated freeze and hypoxia exposures in the summer. Ratios were obtained by normalizing the MO_2_ after exposure to the baseline. A ratio above 1 (grey dotted line) generally signifies an increase in MO_2_ and a ratio below 1 generally signifies a decrease in MO_2_. n=7 for all experiments in the winter. In the summer, n=6 for single exposures to freezing and hypoxia, and n=7 for repeated exposures to freezing and hypoxia. Teal triangles represent mean MO_2_ after freezing and black circles represent mean MO_2_ after hypoxia exposure. Error bars signify the standard error of the mean.

To better understand the duration of the metabolic perturbation following freezing, we examined how MO_2_ varied after a period of recovery. MO_2_ was measured 24 hours after the final exposure for single and repeated exposures to freezing and immersion in the winter and the summer. To compare how MO_2_ changed from the baseline after treatment, the mean over the 40 minute period compared using a repeated measures ANOVA with timepoint as the fixed effect and the individual as the random effect. In the winter, we found that there was a statistically significant increase after a single hypoxia exposure, but a significant decrease after repeated hypoxia exposures. There was a significant increase in MO_2_ ratio 24 hours after a single freeze in the summer. Statistical details are reported in Table S3.

Similarly, MO_2_ ratios were used to account for individual variation, and we calculated each individual’s MO_2_ ratio after normalizing mussel MO_2_ after exposure to their initial pre-exposure baseline. Using a repeated measures ANOVA with the individual as the random effect, we compared the ratios immediately after exposure (i.e. no recovery) and after 24 hours of recovery. Mussels undergoing a single freezing exposure in the winter showed a significantly increased MO_2_ ratio (*χ*^2^=11.07, df=1, p<0.001) after 24 hours while there was no significant change for a single hypoxia exposure (*χ*^2^=0.0087, df=1, p=0.93). For repeated exposures in the winter, there is no significant change in MO_2_ ratio after freezing (*χ*^2^=1.06, df=1, p=0.30) nor hypoxia exposures (*χ*^2^=0.54, df=1, p=0.46) after 24 hours.

However, we found that there was a significant increase in the MO_2_ ratio 24 hours after a single freeze exposure in the summer (*χ*^2^=8.10, df=1, p<0.01) and a significant decrease 24 hours after a single hypoxia exposure (*χ*^2^=8.36, df=1, p<0.01). There was no significant change 24 hours after repeated freezing (*χ*^2^=0.80, df=1, p=0.37) and similarly 24 hours after repeated hypoxia exposures (*χ*^2^=0.018, df=1, p=0.89).

## Discussion

Intertidal mussels in temperate and polar regions can experience freezing naturally during low tides, therefore we examined changes to aerobic metabolism after single and repeated freezes and across seasons. We hypothesized that freezing in the winter and being frozen repeatedly would induce less metabolic perturbation than freezing only once or in the summer. We found that immediately after thawing from a single freeze, aerobic metabolism was depressed in mussels but recovered within 24 hours and we found that exposure to repeated freezes in the winter appeared to cause the least overall metabolic perturbation to mussels. This variation suggests differences in the underlying physiological consequences of freezing across both freezing frequency and season.

### Effects of single freezes

One of the most consistent patterns we observed across all seasons is that aerobic metabolism was depressed immediately after a single freeze-thaw. There are two potential explanations for this depression. Either this may represent an active metabolic suppression, or it may be indicative of damage to the oxygen cascade. Metabolic suppression has been observed in other intertidal organisms as a potentially adaptive mechanism against thermal stress (Hui et al., 2020). A decrease in metabolic rate has also been observed in terrestrial invertebrates immediately after thawing (Sinclair et al., 2004, 2013) and has been suggested as a mechanism to circumvent ischemia-reperfusion damage (Joanisse & Storey, 1998). Alternatively, this depression may be indicative of damage to the oxygen cascade. For example, damage to the gills implicated in oxygen uptake were observed after a freezing event in *M. trossulus* (Kennedy, 2022). This may impair oxygen uptake, and therefore a decrease in MO_2_ may be indicative of damage to the capacity to take up oxygen. As mussels are more freeze tolerant in the winter (Kennedy et al., 2020), mussels may be less susceptible to freezing damage in this season. Therefore, if the metabolic depression was driven by damage, differences in the metabolic depression due to damage would be expected. However, we found that aerobic metabolism was depressed to a similar magnitude across seasons despite the expected differences in seasonal freeze tolerance. This suggests that damage to the oxygen cascade may be a lesser contributor to the observed metabolic depression and may instead be driven by suppressive mechanisms. Additionally, after 24 hours of recovery, we found that metabolic rates were no longer depressed regardless of season, indicating that the initial depression is transient and that this may represent the cessation of a metabolic suppression. The potential mechanisms of metabolic suppression after cold stress in intertidal bivalves have yet to be explored.

Additionally, we tested whether the metabolic depression is due to ice formation or low temperature exposure. Mussels which froze at −5.5 °C tended to have metabolic rates resembling mussels which were frozen at a lower temperature (−6.2 °C). By contrast, the metabolic rate of mussels which did not freeze at −5.5 °C (i.e., emersed at low temperatures) more closely resembled the metabolic rate of a hypoxia exposure at a higher temperature (15 °C). Interestingly some individuals also displayed lower oxygen consumption rates after only a cold exposure, suggesting that low temperatures may be a sufficient trigger for depressing aerobic metabolism. However, the lower amount of variation in aerobic metabolism after freezing suggests that while this response can be triggered by chilling and may depend on the individual, freezing more consistently exceeds the threshold to trigger the metabolic depression in all individuals. These results are similar to what was observed in the Antarctic insects *Hydromedion sparsutum* and *Perimylops antarcticus*, where chilling non-significantly depressed aerobic metabolism (Block et al., 1998). However, they only noted an increase in aerobic metabolism after thawing, suggesting that there are distinct physiological differences between chilling and freezing. Additionally, it is possible that a metabolic elevation expected after hypoxia exposure was not observed here due to lower accumulation of oxygen debt at lower temperatures (Hicks & McMahon, 2002). Taken together, we conclude that the metabolic patterns observed after single freezes may be better attributed to ice crystal formation associated with freezing and not solely by the low temperature exposure.

### Effects of repeated freezes

Unlike single freezes, the metabolic response immediately after repeated freezes varied across seasons. Similar to single freezes, in the fall, aerobic metabolism was depressed after repeated freeze-thaws. In the winter, aerobic metabolism after repeated freeze-thaw was unchanged from the baseline and in the summer, repeated freeze-thaw led to an elevation in aerobic metabolism. After 24 hours of recovery in the winter, there was a slight, but non-significant, increase in oxygen consumption after repeated freezing, whereas in the summer, aerobic metabolism was immediately elevated following freezing and was maintained for at least 24h. In the fall, the metabolic depression in mussels after repeated freezes may share similar biochemical underpinnings as single freezes. However, the lack of a metabolic depression in the winter and the summer may relate to the activity occurring during the periods of repair between each freeze-thaw cycle, and the elevation in metabolic rate may be attributed to several possible reasons. For example, HSP70 induction occurs during these periods and is energetically costly and may have helped counteract the freezing stress and circumvent the period of metabolic depression (Gill et al., 2023). Additionally, ATP is consumed during freezing (Storey & Churchill, 1995) and the regeneration of ATP represents another metabolic cost. Alternatively, elevated metabolic rates can also be indicative of damage. For instance, damage to the mitochondrial membrane may increase proton leak, and therefore MO_2_ may be increased as a compensatory response when respiring aerobically (Štětina et al., 2020; Štětina & Koštál, 2024). Cellular or tissue repair is also likely to be energetically costly and consume ATP. Currently, there is one other study which has directly examined whole organism metabolic rate after repeated freezing in *Pyrrharctia isabella* and it is noted that there is a non-significant trend towards a depressed metabolism 24 hours after repeated freezing (Marshall & Sinclair, 2011). While there is still evidence of damage observed at this time point, the lowered metabolic rate may instead represent different strategies used by insects to survive freezing which may involve repair at a later time point compared to intertidal organisms which must embrace the opportunity to aerobically respire and also feed at each high tide.

The difference in response to repeated freezes across seasons may relate to the amount of stress induced by freezing due to plasticity in freeze tolerance across seasons (Kennedy et al., 2020). As such, there may have been differences in the physiological stress incurred from freezing, but also potentially a difference in the effectiveness of repair processes in between each freeze. There is also more variation in the response after repeated freezes when compared to single freezes. Since repeated freezes are less lethal compared to a single prolonged freeze, this overall milder treatment may also in part drive the variability in the response as opposed to single freezes which did not display a clear seasonal effect (Gill et al., 2023). It is possible that a certain level of freezing stress needs to be accumulated to elicit a metabolic suppression and eventual restoration to the baseline.

### Effects of hypoxia on the response to freezing

Freezing can physiologically be considered a combination of the stress of ice crystal formation and hypoxia stress as these events not only co-occur in the intertidal, but ischemia occurs during freezing (Storey and Churchill, 1995). Therefore, we compared the responses between single freezing and hypoxia exposures. Metabolic rates are found to increase after mussels were emersed for 6 hours and this response is consistent across seasons. During emersion, mussels are largely hypoxic and therefore oxygen debt is accrued. Therefore, following immersion, aerobic metabolism was elevated likely indicating that oxygen debt was being repaid (Ellington, 1983). Since the response observed after a hypoxia exposure is opposite to that of a freezing exposure, it is unlikely that the metabolic response after freezing is driven by hypoxia stress but rather by freezing stress.

We also compared the metabolic responses between repeated freezing and hypoxia exposures. In the fall, mussels showed a depression after repeated freezing but an elevation to repeated hypoxia exposures. This similarly suggests that the metabolic response to freezing is not driven by hypoxia stress. However, in the winter and the summer, mussels initially presented a similar metabolic response to freezing and emersion in the winter and the summer. However, after 24 hours, the metabolic response becomes more distinct from the response of emersion, similarly indicating that the effect of emersion is physiologically different from that of freezing. In both the winter and the summer, there is no elevation in metabolic rate after hypoxia exposure, potentially indicating that due to a relatively short period of time spent emersed and also due to the recovery periods, there is minimal accumulation of oxygen debt. Therefore, the metabolic consequences of freezing are more likely driven by the freezing and not of the hypoxia stress that co-occurs with freezing in the intertidal.

This series of experiments only capture aerobic metabolism despite the presence of numerous anaerobic pathways which may be activated during freezing. Freezing induces ischemia as the hemolymph is frozen and therefore gas exchange with tissues is effectively halted. A previous study examining anaerobic byproduct accumulation after freezing shows that there is no significant difference between frozen and unfrozen mussels after 24 hours of recovery (Kennedy et al., 2020). However, it is possible that the byproducts are cleared at that time point, and the timepoint immediately after thawing should also be examined to see whether significant anaerobic activity occurs during freezing before switching to aerobic pathways upon emersion in seawater.

## Conclusion

This study demonstrates the metabolic responses associated with ecologically-relevant freeze-thaws at the whole organism level. Freezing singly generally leads to a transient depression in metabolism, followed by an increase in aerobic metabolism which is also generally observed after repeated freezing events. These responses may represent adaptive mechanisms which aid in the freeze tolerance of intertidal mussels. Despite a warming world, freezing still plays a relevant threat with the predicted increase in frequency of extreme weather events (Francis & Vavrus, 2012). Better understanding the energetics of freezing may help us better understand how the intertidal ecosystem may be affected. While this study uncovers seasonal differences in the response to freezing, specific mechanisms which govern the seasonal differences to freeze tolerance are unknown. To better understand these fundamental mechanisms of intertidal freeze tolerance, we propose that future work should explore direct evidence for freezing damage and repair processes.

## Acknowledgements

KEM is supported by an NSERC Discovery Grant and JCCY was supported by an NSERC USRA and CGS-M during this work. The authors thank Dr. Patricia Schulte and Dr. Jeffrey Richards for their helpful suggestions on this project.

