## Supplemental data for "Energetic consequences of single and repeated freezing in the intertidal mussel, *Mytilus trossulus*"

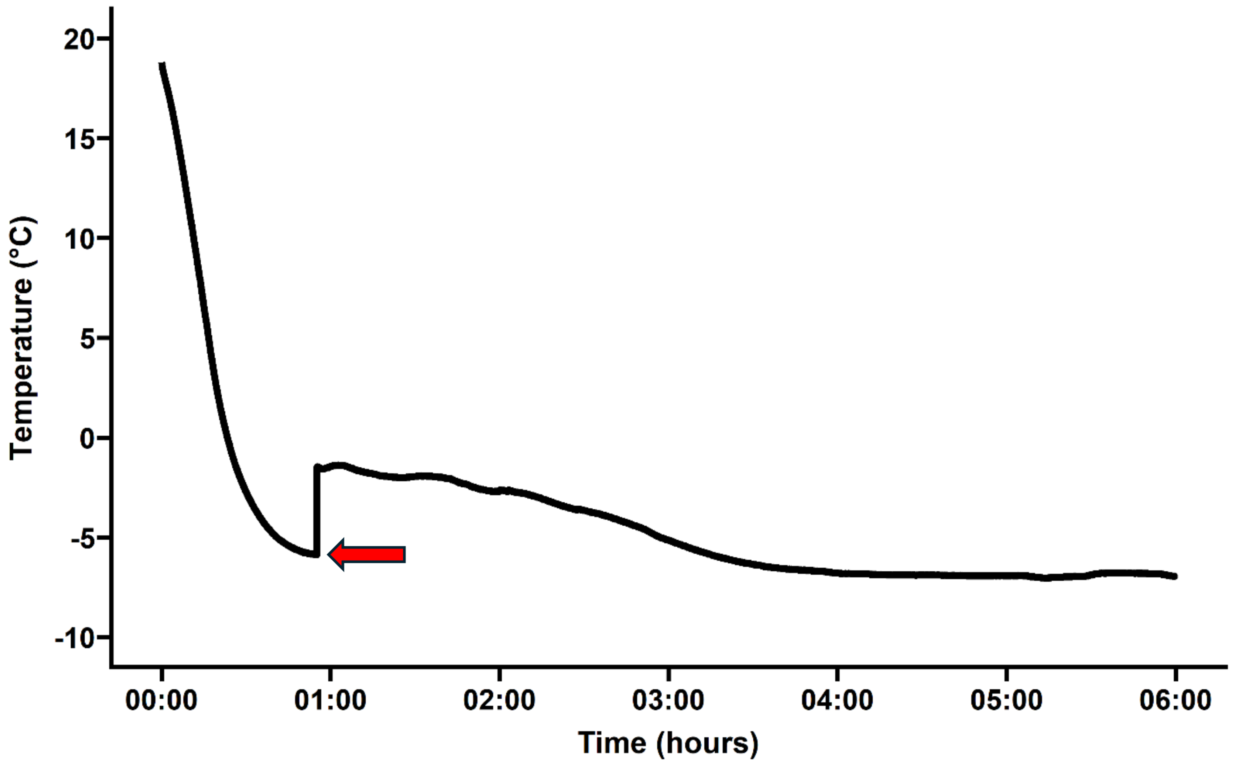


**Figure S1. Example temperature trace of a mussel during a 6 hour cold exposure.** Body temperature of a mussel was tracked using a thermocouple attached externally to the shell. The sharp increase in body temperature (red arrow) due to the freezing exotherm indicates a freezing event.

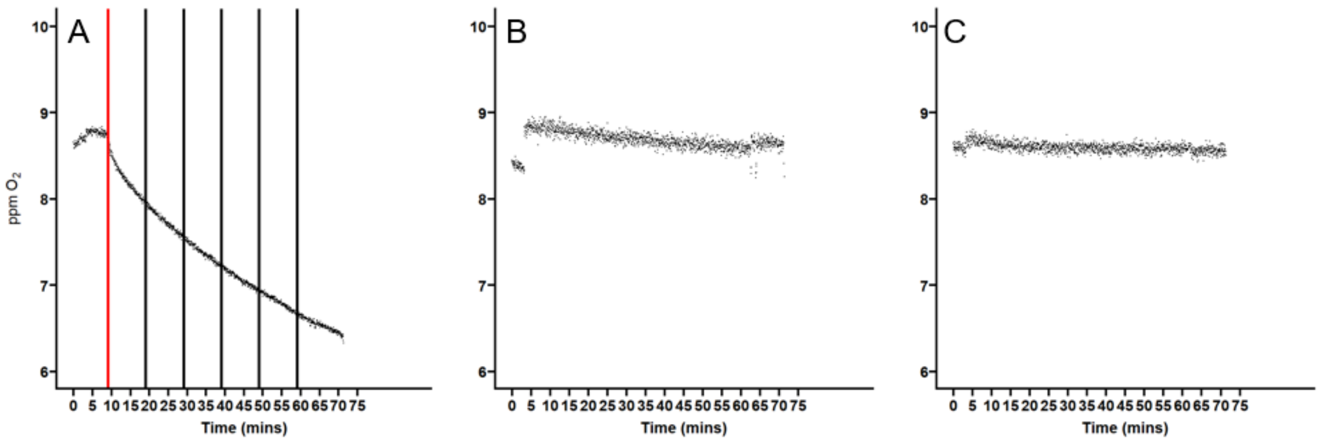


**Figure S2. Sample oxygen content traces recorded by oxygen sensor spots from a respirometer chamber with a mussel in seawater.** (A) shows a trace with a detectable signal from a mussel and (B) shows a trace with no detectable signal from a mussel (C) represents the background measurement (seawater only). On panel A, the red line indicates the moment of detectable oxygen consumption, and the sections between black lines indicate the four 10 minute periods used to calculate the oxygen consumption rate.

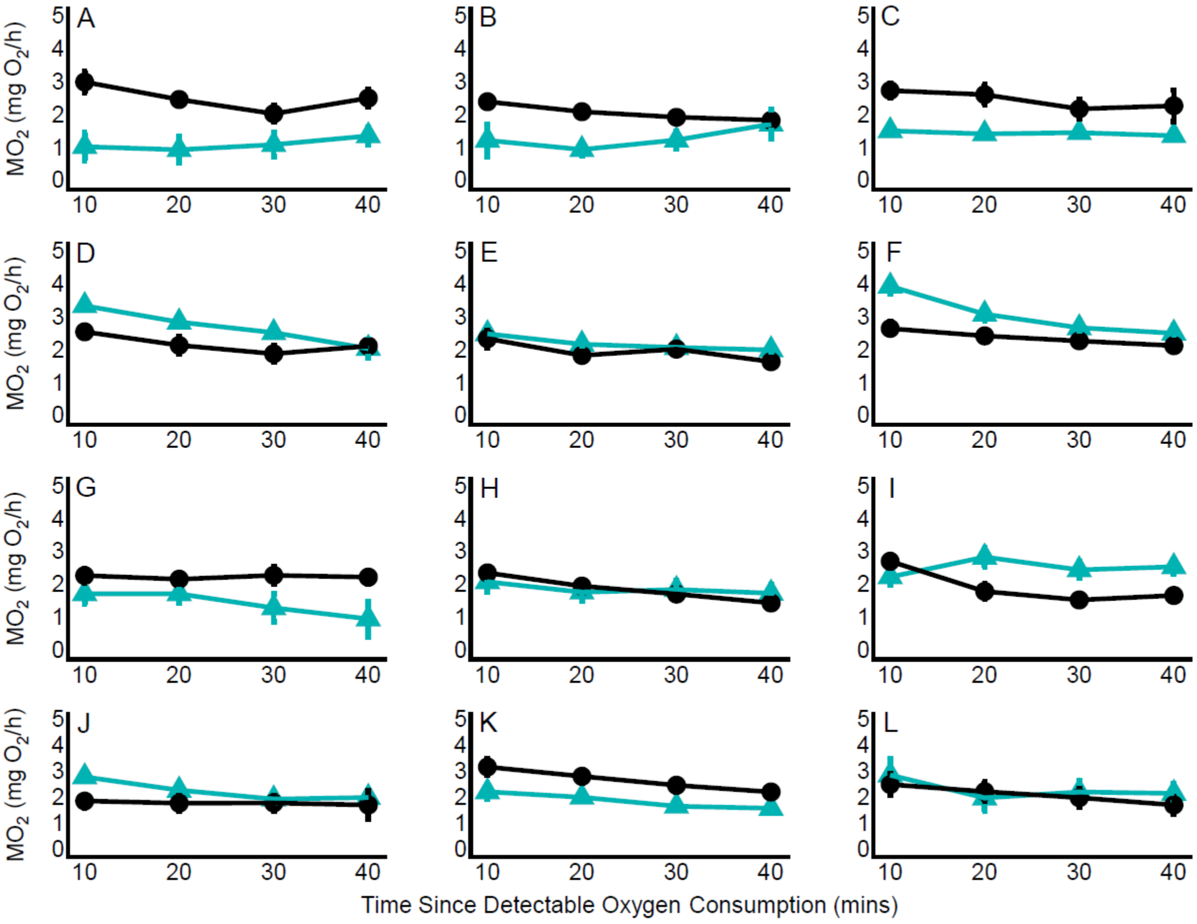


**Figure S3. Comparison of $MO_2$ before and after freezing or hypoxia exposure across seasons**

Comparison of $MO_2$ at 15 °C of *M. trossulus* across before (black) and after (blue) experimental exposures. Data represent a 40 minute period after detectable oxygen consumption in the respirometry chamber. Comparisons are shown for single freezes in the fall, winter, and summer (A, B, C, respectively), for single hypoxia exposures in the fall, winter, and summer (D, E, F, respectively), for repeated freezes in the fall, winter, and summer (G, H, I, respectively), and for repeated hypoxia exposures in the fall, winter, and summer (J, K, L, respectively). n=7 for all experiments except for single and repeated freezes in the summer. Teal triangles represent pre-exposure values while black circles represent post-exposure values. Error bars represent the standard error of the mean.

**Table S1. Summary of environmental measurements on sampling days and experimental exposures.** Tidal data was sourced from a station situated at Point Grey, Vancouver, B.C. Canada.

| Collection Date | Site Salinity (ppt) | Site air Temperature (°C) | Site water temperature (°C) | Low tide | Experiment Dates | Experiment |
|---|---|---|---|---|---|---|
| Sept 8, 2022 | 19.5 | 18.3 | 19.3 | 0.8m (10:34am) | Sept 13, 2022<br>Sept 15, 2022<br>Sept 19 2022 | Single 6h Freeze (-5.5 °C)<br>Single 6h Freeze (-6.2 °C)<br>Single 6h Hypoxia (15 °C) |
| Sept 22, 2022 | 19.6 | 15.5 | 14.1 | 1.4m (10:10am) | Sept 26-28, 2022<br>Sept 26-28, 2022<br>Sept 30, 2022 | Repeated Freeze (-6.2 °C)<br>Repeated Hypoxia (15°C)<br>Single Freeze (-5.5 °C) |
| Jan 23, 2023 | 25.4 | 6.5 | 7.1 | 0.1m, (12:10am) | Jan 25-27, 2023<br>Jan 30-Feb 1, 2023 | Repeated Freeze (-6.2 °C)<br>Repeated Hypoxia (15 °C) |
| Feb 3, 2023 | 25 | 13.5 | 6.9 | 0.8m (10:38pm) | February 6, 2023<br>February 6, 2023 | Single 6h Freeze (-6.2 °C)<br>Single 6h Hypoxia (15 °C) |
| Aug 16, 2023 | 21.5 | 23.2 | 20.9 | 0.8m (12:36pm) | August 19, 2023<br>August 20, 2023 | Single 6h freeze (-6.2 °C)<br>Single 6h hypoxia (15 °C) |
| Aug 30, 2023 | 18.2 | 21.4 | 22.4 | 0.6m (11:55pm) | September 1-3, 2023 | Repeated Freeze (-6.2 °C)<br>Repeated Hypoxia (15 °C) |

**Table S2. Summary statistics for comparison of $MO_2$ before and after freezing or hypoxia exposure across seasons**

| Treatment | Season | $\chi^2$ | df | p | Figure |
|---|---|---|---|---|---|
| Single freeze | Fall | 15.53 | 1 | <0.001 | A |
| | Winter | 12.89 | 1 | <0.001 | B |
| | Summer | 16.42 | 1 | <0.001 | C |
| Single Hypoxia | Fall | 21.71 | 1 | <0.001 | D |
| | Winter | 8.40 | 1 | <0.01 | E |
| | Summer | 19.09 | 1 | <0.001 | F |
| Repeated Freeze | Fall | 5.40 | 1 | <0.05 | G |
| | Winter | 0.0006 | 1 | 0.980 | H |
| | Summer | 5.28 | 1 | <0.05 | I |
| Repeated Hypoxia | Fall | 6.50 | 1 | <0.05 | J |
| | Winter | 38.00 | 1 | <0.001 | K |
| | Summer | 0.67 | 1 | 0.412 | L |

**Table S3. Summary statistics for comparisons of the mean $MO_2$ of *M. trossulus* between their initial baseline $MO_2$ and 24 hours after single or repeated freeze or hypoxia exposures.**

| | Treatment | $\chi^2$ | df | p |
|---|---|---|---|---|
| Winter | Single Freeze | 3.64 | 1 | 0.056 |
| | Single Hypoxia | 11.06 | 1 | <0.001 |
| | Repeated Freeze | 2.72 | 1 | 0.098 |
| | Repeated Hypoxia | 28.35 | 1 | <0.001 |
| Summer | Single Freeze | 0.11 | 1 | 0.74 |
| | Single Hypoxia | 2.84 | 1 | 0.092 |
| | Repeated Freeze | 4.86 | 1 | 0.027 |
| | Repeated Hypoxia | 0.17 | 1 | 0.68 |
